# DeGlyPHER: an ultrasensitive method for analysis of viral spike N-glycoforms

**DOI:** 10.1101/2021.05.13.444041

**Authors:** Sabyasachi Baboo, Jolene K Diedrich, Salvador Martínez-Bartolomé, Xiaoning Wang, Torben Schiffner, Bettina Groschel, William R Schief, James C Paulson, John R Yates

## Abstract

Viruses can evade the host immune system by displaying numerous glycans on their surface “spike-proteins” that cover immune epitopes. We have developed an ultrasensitive “single pot” method to assess glycan occupancy and the extent of glycan processing from high-mannose to complex forms at each N-glycosylation site. Though aimed at characterizing glycosylation of viral spike-proteins as potential vaccines, this method is applicable for analysis of site-specific glycosylation of any glycoprotein.

## Introduction

Viral spike-proteins initiate virus entry into host cells and are the primary targets of vaccine design. Spike-proteins are often heavily N-glycosylated which help to shield the protein from the host immune response^1^. These glycans add complexity to the production and characterization of recombinant protein-based vaccines^2^. This is particularly a concern for characterization of the envelope spike-protein (*Env*) of the Human Immunodeficiency Virus (HIV) comprising a trimer with each monomer containing 26-30 unique N-linked glycosylation sites (NGS) as defined by the sequon `NX[S|T]`, where `X` is any amino acid except `P`^3^. To address this analytical challenge several mass spectrometry-based strategies using multiple proteases^4–6^have been implemented to create sufficient numbers of peptides unique to each glycosylation site^3,7,8^. In these strategies, individual aliquots are digested with each protease and analyzed separately by liquid chromatography-mass spectrometry (LC-MS/MS), or pooled and analyzed together. To broadly characterize the nature of the glycosylation at each NGS, we had previously introduced the use of endoglycosidases that create residual mass signatures^3^. This helped us to determine the degree of glycan occupancy, and the degree of glycan processing – from the high mannose form that is initially attached to the protein, which may mature into the complex form when mannose residues are replaced by “terminal” monosaccharide sequences. To achieve coverage for all NGS, we had combined several digestions performed with different proteases and achieved >95% sequence coverage. Here we show that it is possible to replace these multiple proteolytic digestions with a single Proteinase K (PK) digestion, and moreover through careful choice of volatile buffers we have developed an improved “single pot” strategy with significantly increased sensitivity. We name this strategy to analyze glycoforms as DeGlyPHER (*De*glycosylation-dependent *Gly*can/*P*roteomic *H*eterogeneity *E*valuation *R*eport).

## Results and Discussion

PK is a broadly specific serine-protease that has previously been exploited for its potential to generate overlapping peptides and high sequence coverage^9^. The redundancy afforded by overlapping sequences significantly increases confidence in identifications, especially when covalent modifications are present^4^. However, because proteinase K is an aggressive protease it is necessary to attenuate its proteolytic activity to obtain high sequence coverage of proteins.

Attenuation of PK can be achieved using suboptimal reaction conditions^9^ (to reduce the rate of enzyme activity) and limited reaction time. Using a mildly acidic, chaotrope-free solution to attenuate the activity of PK, we were able to achieve >95% sequence coverage of candidate viral spike-proteins and identify all NGS. The proteomic strategy of DeGlyPHER is conceptually similar to our previous approach^3^, but it is significantly faster and more sensitive. These improvements result from three key changes to the strategy: [1] using only mass spectrometry-compatible constituents, samples are processed in a single solution except the final step of PNGase F deglycosylation; [2] reaction volumes are kept to a minimum (5-8 μl) to increase the rate of reaction to limit sample loss on surfaces and minimize freeze-drying time; and [3] the use of PK provides faster digestion and excellent sequence and NGS coverage with less starting material. DeGlyPHER reduces sample preparation time from 3 days to 1 day, reduces LC-MS/MS run times by 9 to 24-fold, and can achieve 90-180 times higher sensitivity than existing methods^3,7,8^. In addition, we have developed the data analysis tool GlycoMSQuant which further reduces analysis times and simplifies data analysis.

We tested DeGlyPHER on BG505 SOSIP.664 MD39^10^, a stabilized native-like HIV *Env* trimer being developed for an HIV vaccination strategy^11^ targeting germline precursors of broadly neutralizing antibodies (bNAbs) that are impacted by N-glycans. As in our previous approach^3^, the glycosylated peptides generated by PK digestion were sequentially deglycosylated; first with Endo H to remove high-mannose and hybrid N-glycans, and then with PNGase F, which removes all remaining N-glycans. The resulting residual masses on asparagine (`N`) in NGS is +203 Da at sites occupied by high-mannose/hybrid N-glycans or +3 Da at sites occupied by complex N-glycans when PNGase F deglycosylation is carried out in the presence of H_2_^18^O (differentiating these sites from any deamidated `N`s). Unoccupied NGS results in no (+0 Da) residual mass on `N`. Using DeGlyPHER (***Fig. 1a***), we achieved >99% amino acid sequence coverage and identified all theoretically possible 27 NGSs from a single LC-MS/MS run of 0.5 μg of peptides generated from a starting material of 5 μg purified protein (***Fig. 1b*** and ***Supplementary Figures 1a,b***). We used semi-quantitative label-free analysis based on precursor peak areas to calculate the proportion of N-glycan occupancy (unoccupied: complex: high-mannose/hybrid N-glycans) for each NGS. We reanalyzed the N-glycan microheterogeneity pattern on BG505 SOSIP.664^12^ HIV *Env* trimer from data obtained using our previous approach^3^ and compared it with results using DeGlyPHER (***Supplementary Figures 2***). The results with DeGlyPHER were highly comparable to those using our original approach, in spite of being processed differently and the samples being prepared at different times in different laboratories.

**Figure 1.**
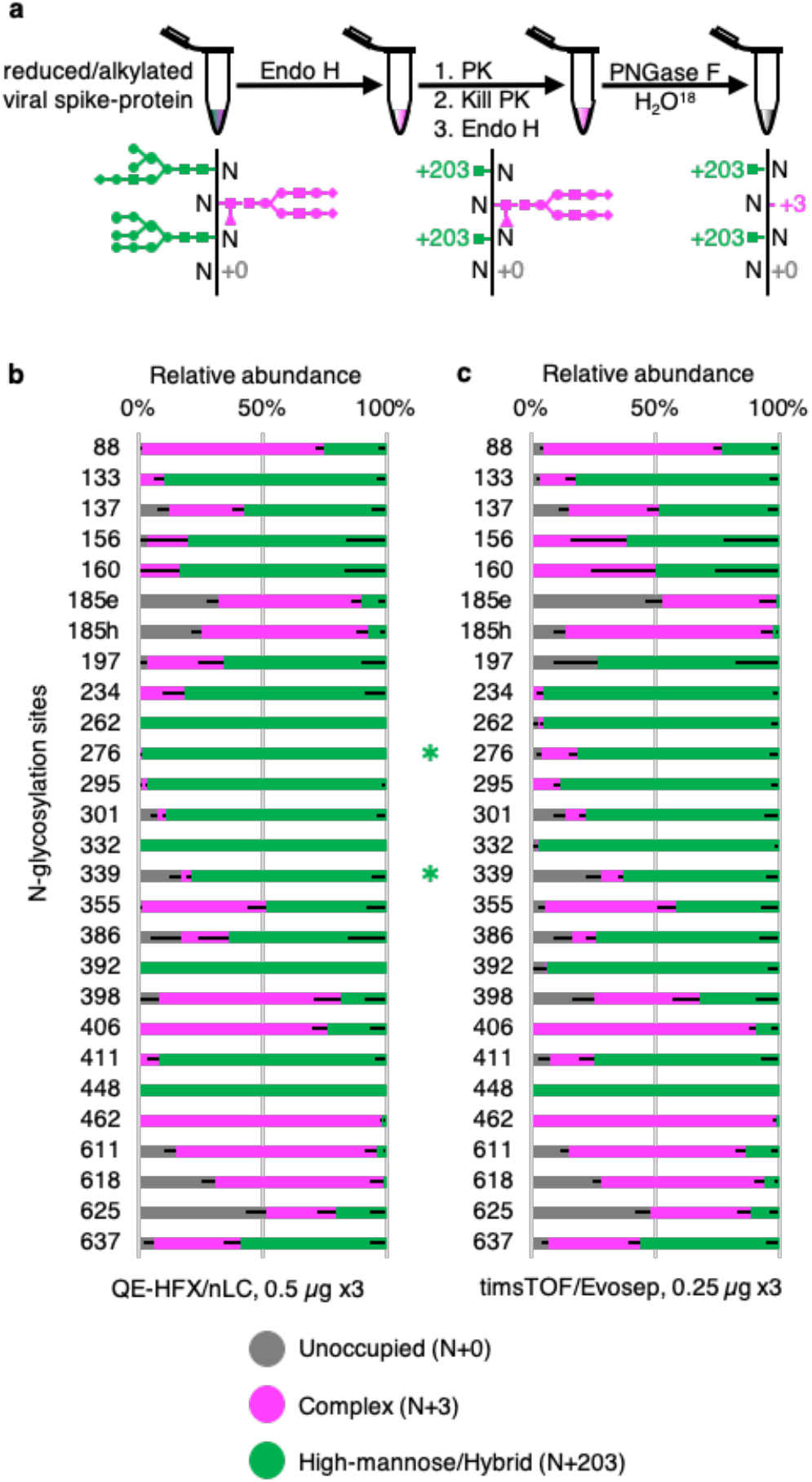
Proteinase K and glycosidase treatment provides N-glycan microheterogeneity in BG505 SOSIP.664 MD39 trimer. (**a**) DeGlyPHER workflow, resulting in peptides with expected mass modifications at NGS representing N-glycosylation states. (**b**) Pattern observed using 4 h triplicates on an QE-HFX/nLC platform. (**c**) Pattern observed using 88 min triplicates on timsTOF/Evosep platform. N-glycosylation states are color-coded. Error bars represent mean-SEM. Between (**b**) and (**c**), the pattern is similar, and any significant difference (BH-corrected *p*-value <0.05) in proportion of a certain N-glycosylation state at an NGS is represented by color-coded **\***.

Initial results demonstrated that DeGlyPHER is at least 18 times more sensitive than our previous approach^3^ even though it uses a simpler and shorter workflow. To evaluate the limit of sensitivity of DeGlyPHER, we processed progressively decreasing amounts of starting material, ranging from 1 μg to 5 ng. We observed that a single LC-MS/MS run with 1 μg of starting material was enough to cover >95% of the amino acid sequence and all NGS (***Fig. 2a***), which is 90 times more sensitive than our previous approach^3^. Major differences in microheterogeneity at each NGS were generally observed when we started with <100 ng material (***Supplementary Figure 1c***). This is likely due to low sampling as evidenced by a decrease in amino acid sequence and NGS coverage (***Fig. 2a***), as well as the absolute number of identified peptides representing each NGS (***Supplementary Figure 1d***).

**Figure 2.**
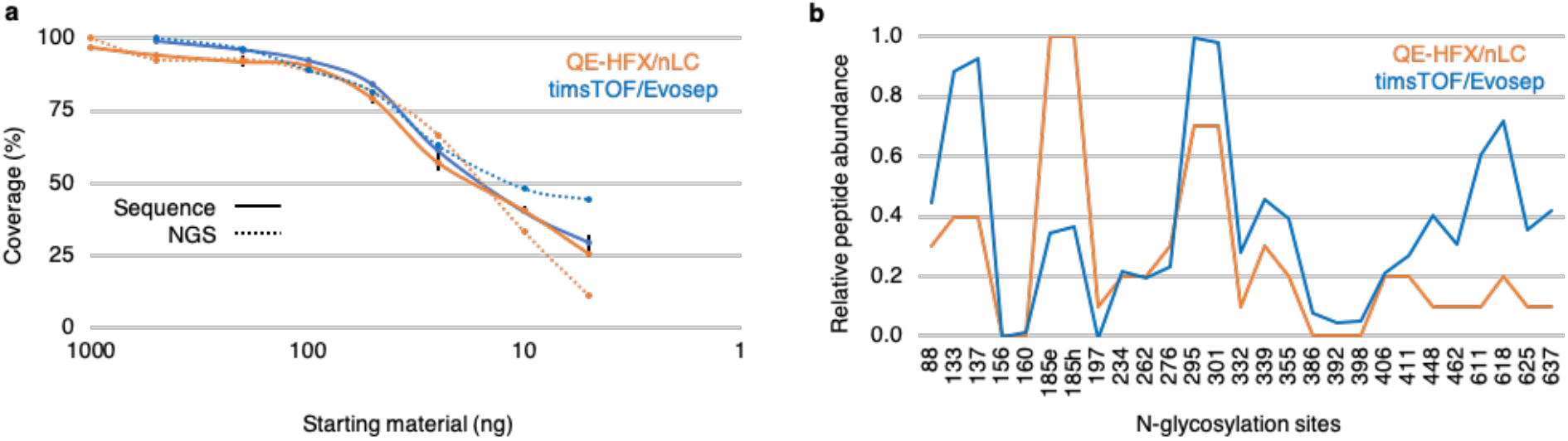
Factors affecting limit of sensitivity. (**a**) Decreasing trend of sequence and NGS coverage observed as starting material is diluted 200-fold (1 μg to 5 ng, QE-HFX/nLC) or 100-fold (0.5 μg to 5 ng, timsTOF/Evosep), using triplicates for each amount of starting material. The limit of sensitivity is revealed. (**b**) Relative abundance of peptides identified per NGS across the dilution series for the 2 LC-MS/MS platforms used. This comparison reveals a non-uniform digestion pattern that may be attributed to steric hindrance offered by the glycoprotein or characteristic behavior of individual peptides in LC-MS/MS. Error bars represent mean±SEM. Values for sequence and NGS coverage are mean

DeGlyPHER is agnostic to mass spectrometry platform (***Fig. 2b***). A timsTOF Pro mass-spectrometer coupled to an Evosep One HPLC (timsTOF/Evosep)^13^ was used to achieve >99% sequence coverage and identification of all NGS using a single LC-MS/MS run with 0.5 μg of starting material and an 88-minute LC gradient (***Fig. 1c*** and ***Supplementary Figures 3a,b***). Thus, the sensitivity of DeGlyPHER on this platform was 180 times higher than our previous approach^3^.

N-glycan heterogeneity reflects the immunogenicity of the viral spike-protein and is critical for designing vaccines^14^. The reproducibility of N-glycan heterogeneity patterns obtained with DeGlyPHER suggests that this is a robust procedure. Variability in sequence coverage is not observed until the limits of detection are reached on an LC-MS/MS platform. Although our results within the same LC-MS platform were reproducible (except some variations when using different proteases, ***Supplementary Figure 5***), we may infer the effects of sampling differences when comparing two different LC-MS platforms (QE-HFX *vs.* timsTOF/Evosep), as in case of `N`156, `N`160, `N`197, `N`386, `N`392 (relative peptide abundance is persistently low) and `N`88, `N`295, `N`301, `N`332, `N`355, `N`406, `N`411 (possible skewing of timsTOF/Evosep identification against `N`+203 peptides when peptide sampling per NGS decreases due to less starting material) (***Fig. 2b*** and ***Supplementary Figures 1c,d*** and ***3c,d***). When enough sampling per NGS is achieved, these variations are diminished (***Figs. 1b,c***).

We attribute improvements in DeGlyPHER to efficient sample handling strategies. We observed reduced sequence coverage if the sample was from the digestion of a small amount of starting material rather than an equal aliquot from a larger sample (***Supplementary Figure 3e***). We infer that the sensitivity differences are not occurring during LC-MS/MS, but that sample is being lost to the reaction-tube surface (during reaction and lyophilization) and the proportion of loss is more pronounced when we start with less material. The kinetics of the enzyme-substrate reaction may also account for sensitivity differences since a more “crowded” reactant environment (low reaction volumes) is expected to result in better reaction kinetics^15^.

The simplicity and high reproducibility of DeGlyPHER will allow for high-throughput analyses of viral spike-proteins and for any glycoprotein whether produced recombinantly or purified from natural sources. Results were highly comparable in the two LC-MS/MS platforms we used. A single analysis using a QE-HFX/nLC platform can determine the complete N-glycan heterogeneity pattern from 1 microgram of purified viral spike-protein. The timsTOF/Evosep platform was observed to be more sensitive than QE-HFX/nLC across all NGS, although limited sampling may not allow us to confidently infer N-glycan microheterogeneity at all NGS. We view the high sensitivity of DeGlyPHER to be an important step to analyze glycosylation of more complex samples such as whole virus or virus in infected blood^16^.

## Methods

### Expression and purification of HIV *Env* trimers

BG505 SOSIP.664^12^ and BG505 SOSIP.664 MD39^10^ *Env* trimers were expressed and purified essentially as described previously^10^. Briefly, sequences with codons optimized for expression in human cells were synthesized and cloned into pHLSec between *AgeI*/*KpnI* by Genscript. The constructs were co-transfected with Furin-encoding plasmid, using polyethylenimine in Freestyle 293F cells cultured in 293 FreeStyle media (Thermo Fisher Scientific). Where indicated, 15 μM sterile-filtered Kifunensine (Cayman Chemical) was added after transfection. After 6-7 days, supernatant was collected after passing through 0.22 μm filter (Nalgene), and the C-terminally His-tagged trimers were purified using a HisTrap affinity column (Cytiva) with a linear elution gradient from 20-500 mM imidazole, followed by a Superdex 200 Increase SEC column (Cytiva) in Tris-buffered saline/TBS (20 mM Tris, 100 mM NaCl, pH 7.5). The oligomeric state and purity of trimer was verified using size exclusion chromatography coupled with multi-angle light scattering (SEC-MALS; DAWN HELEOS II/ Optilab T-rEX, Wyatt Technology).

### Proteinase K treatment and deglycosylation

HIV Env trimer was exchanged to water using Microcon Ultracel-10 centrifugal device (Millipore Sigma). Trimer was reduced with 5 mM tris(2-carboxyethyl)phosphine hydrochloride (TCEP-HCl, Thermo Scientific) and alkylated with 10 mM 2-chloroacetamide (Sigma Aldrich) in 100 mM ammonium acetate for 20 min at room temperature (RT, 24°C). Initial protein-level deglycosylation was performed using 250 U of Endo H (New England Biolabs) for up to 5 μg trimer, for 1 h at 37°C (pH 5.5-6.0). Trimer was digested with 1:25 Proteinase K (Sigma Aldrich) for 4 h at 37°C (pH 5.5-6.0). PK was denatured by incubating at 90°C for 15 min, then cooled to RT. Peptides were deglycosylated again with 250 U Endo H for 1 h at 37°C (pH 5.5-6.0), then frozen at –80°C and lyophilized. 100 U PNGase F (New England Biolabs) was lyophilized (for up to 5 μg trimer), resuspended in 100 mM ammonium bicarbonate prepared in H_2_^18^O (97% ^18^O, Sigma-Aldrich), and added to the lyophilized peptides. The resulting 5-8 μl reaction solutions (except PNGase F reaction, in 5-20 μl) were then incubated for 1 h at 37°C (pH 8.0-8.5) in 0.2 ml PCR tubes on a thermocycler with heated lid.

### Validating efficiency of glycosidases

BG505 SOSIP.664 HIV *Env* trimer glycosylated with only high-mannose N-glycans (purified from cells treated with Kifunensine^17^, which inhibits processing of high-mannose N-glycans to complex N-glycans during protein maturation), was sequentially treated with Endo H, followed by PNGase F. After treatment with both enzymes, 99.2% of identified peptides were `N`+203; 100% of peptides identified with only PNGase F treatment were `N`+3 (***Supplementary Figures 4a,b***; proportions do not consider unoccupied NGS because they remained similar in both experiments – 8-10%). We realize the possibility that glycosidase PNGase F may occasionally cleave the remnant GlcNAc (N-Acetylglucosamine)^18^ post-Endo H processing of high-mannose/hybrid N-glycans and thus, convert the mass modification characteristic of high-mannose/hybrid (+203) to complex (+3) N-glycans, which would affect our analyses. However, we have not observed any significant evidence of this possibility in our results, though it may explain why we observe a few peptides with +3 mass modified NGS in Kifunensine treated samples (***Supplementary Figure 4a***).

### Trypsin proteolysis

The Proteinase K/deglycosylation method described above was followed, except PK was replaced with trypsin and reactions were incubated overnight at 37°C. Trypsin generated a lower total number of peptides than PK, but we obtained >95% sequence coverage, including 26 of 27 NGS (***Supplementary Figure 5a***). Variations in N-glycan microheterogeneity at certain NGS may be explained by low sampling at these sites (`N`386, `N`392) or difference in cleavage-specificity between PK and trypsin (`N`88, `N`611) (***Supplementary Figures 5b,c***).

### LC-MS/MS

#### Q Exactive HF-X with EASY-nLC 1200

Samples were analyzed on an Q Exactive HF-X mass spectrometer (Thermo). Samples were injected directly onto a 25 cm, 100 μm ID column packed with BEH 1.7 μm C18 resin (Waters). Samples were separated at a flow rate of 300 nL/min on an EASY-nLC 1200 (Thermo). Buffers A and B were 0.1% formic acid in 5% and 80% acetonitrile, respectively. The following gradient was used: 1–25% B over 160 min, an increase to 40% B over 40 min, an increase to 90% B over another 10 min and 30 min at 90% B for a total run time of 240 min. Column was re-equilibrated with solution A prior to the injection of sample. Peptides were eluted from the tip of the column and nanosprayed directly into the mass spectrometer by application of 2.8 kV at the back of the column. The mass spectrometer was operated in a data dependent mode. Full MS1 scans were collected in the Orbitrap at 120,000 resolution. The ten most abundant ions per scan were selected for HCD MS/MS at 25 NCE. Dynamic exclusion was enabled with exclusion duration of 10 s and singly charged ions were excluded.

#### timsTOF Pro with Evosep One

Samples were loaded onto EvoTips following manufacturer protocol. The samples were run on an Evosep One (Evosep) coupled to a timsTOF Pro (Bruker Daltonics). Samples were separated on a 15 cm × 150 μm ID column with BEH 1.7 μm C18 beads (Waters) and integrated tip pulled in-house using either the 30 SPD or 15 SPD methods. Mobile phases A and B were 0.1% formic acid in water and 0.1% formic acid in acetonitrile, respectively. MS data was acquired in PASEF mode with 1 MS1 survey TIMS-MS and 10 PASEF MS/MS scans acquired per 1.1 s acquisition cycle. Ion accumulation and ramp time in the dual TIMS analyzer was set to 100 ms each and we analyzed the ion mobility range from 1/K_0_ = 0.6 Vs cm^−2^ to 1.6 Vs cm^−2^. Precursor ions for MS/MS analysis were isolated with a 2 Th window for m/z < 700 and 3 Th for m/z >700 with a total m/z range of 100-1700. The collision energy was lowered linearly as a function of increasing mobility starting from 59 eV at 1/K_0_ = 1.6 VS cm^−2^ to 20 eV at 1/K_0_ = 0.6 Vs cm^−2^. Singly charged precursor ions were excluded with a polygon filter, precursors for MS/MS were picked at an intensity threshold of 2,500, target value of 20,000 and with an active exclusion of 24 s.

### Data Processing

Protein and peptide identification were done with Integrated Proteomics Pipeline (IP2, Bruker Scientific LLC). Tandem mass spectra were extracted from raw files using RawConverter^19^ (timstofCoverter for timsTOF Pro data) and searched with ProLuCID^20^ against a database comprising UniProt reviewed (Swiss-Prot) proteome for *Homo sapiens* (UP000005640), UniProt amino acid sequences for Endo H (P04067), PNGase F (Q9XBM8), and Proteinase K (P06873), amino acid sequences for BG505 SOSIP.664^12^ and BG505 SOSIP.664 MD39^10^ (including a preceding secretory signal sequence and followed by 6xHis-tag), and a list of general protein contaminants. The search space included no cleavage-specificity (all fully tryptic and semitryptic peptide candidates for trypsin treatment). Carbamidomethylation (+57.02146 `C`) was considered a static modification. Deamidation in presence of H_2_^18^O (+2.988261 `N`), GlcNAc (+203.079373 `N`), oxidation (+15.994915 `M`) and N-terminal pyroglutamate formation (– 17.026549 `Q`) were considered differential modifications. Data was searched with 50 ppm precursor ion tolerance and 50 ppm fragment ion tolerance. Identified proteins were filtered using DTASelect2^21^ and utilizing a target-decoy database search strategy to limit the false discovery rate to 1%, at the spectrum level^22^. A minimum of 1 peptide per protein and no tryptic end (or 1 tryptic end when treated with trypsin) per peptide were required and precursor delta mass cut-off was fixed at 10 ppm for data acquired with Q Exactive HF-X or 20 ppm for data acquired with timsTOF Pro. Statistical models for peptide mass modification (modstat) were applied (trypstat was additionally applied for trypsin-treated samples). Census2^23^ label-free analysis was performed based on the precursor peak area, with a 10 ppm precursor mass tolerance and 0.1 min retention time tolerance. “Match between runs” was used to find missing peptides between runs for Q Exactive HF-X data (for timsTOF Pro data, reconstructed-MS1 based chromatograms combining isotope peaks for all triggered precursor ions were pre-generated, and then chromatograms were assigned to identified peptides for quantitative analysis, without retrieving missing peptides).

### Data Analysis using GlycoMSQuant

Our new tool GlycoMSQuant v.1.4.1 (https://github.com/proteomicsyates/GlycoMSQuant) was implemented to automate the analysis and to visualize the results. GlycoMSQuant summed precursor peak areas across replicates, discarded peptides without NGS, discarded misidentified peptides when N-glycan remnant-mass modifications were localized to non-NGS asparagines and corrected/fixed N-glycan mislocalization where appropriate. The results were aligned to NGS in *Env* of HXB2^24^ HIV-1 variant.

Precursor peak area was calculated by Census2^23^ from extracted-ion chromatogram (XIC) for each peptide in each replicate. For each NGS (`NX[S|T]`, where `X` is any amino acid except `P`), the “N-glycosylation state” represented by proportions for unoccupied (+0, *u*), complex (+2.988261, *c*) and, high-mannose/hybrid (+203.079373, *h*) N-glycans was calculated as follows.

The sum of the precursor peak areas *S*_*g,pepz*_ was calculated as:

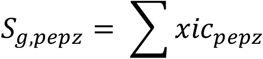

where N-glycosylated peptides with the same sequences and charge were grouped together (*pepz*), *g* is the N-glycosylation state ∈ *G* (*u*, *c*, *h*), and *xic* is the precursor peak area.
For each group (*pepz*), the abundance proportion *%*_*g,pepz*_ of each N-glycosylation state *g* ∈ *G* was calculated as:

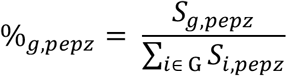

Finally, as each NGS may be covered by multiple groups (*pepz*), the proportion of each N-glycosylation state *g* for a particular NGS (*ngs*) is calculated as the mean of all proportions *%*_*g,pepz*_ of all groups (*pepz*) covering this NGS:

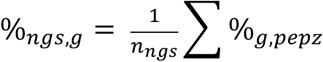

where *n*_*ngs*_ is the number of groups (*pepz*) covering a particular NGS.
The standard error of mean of the proportion of each N-glycosylation state *g ° G* for a particular NGS (*SEM*_*ngs,g*_) was calculated as:

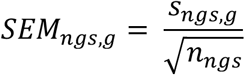

where, *s*_*ngs,g*_ is the standard deviation of *%*_*g,pepz*_ from all groups (*pep*`z`) covering a particular NGS.

Pairwise statistical comparisons of experiments (*a* and *b*) were performed for each *g ∈ G* at each NGS using proportion values *%*_*g,pepz*_ of groups (*pepz*) sharing the NGS, applying the Mann-Whitney U test^25^. Testing *%*_*g,pepz,a*_ *vs. %*_*g,pepz,b*_ individually for *u*, *c* and *h* at each NGS, we calculated *p*-values that were subjected to multiple hypothesis correction using the Benjamini-Hochberg (BH) method^26^. If the corrected *p*-value was <0.05 (for *u*, *c* or *h* at any NGS), then the difference was considered statistically significant.

For ***Supplementary Figure 2***, published data from our previous approach^3^ was reanalyzed using the data analysis workflow described here. Briefly, the data analyzed is from 3 replicates of 3 conditions, each separately analyzed by LC-MS/MS, with total starting material of 90 μg protein, and Census2^23^ label-free analysis was performed simultaneously on all 9 experiments without “match between runs”, and the results analyzed by GlycoMSQuant. This was compared with data obtained from a single LC-MS/MS run (QE-HFX/nLC) with 0.5 μg of peptides generated from a starting material of 5 μg purified protein.

## Acknowledgement

We thank Bruker Daltonics for providing access to timsTOF Pro MS. We thank Bruker Scientific LLC and Robin S Park for making IP2 accessible for data analysis. We thank Titus Jung in J.R.Y. lab for IT support. We thank members of the J.R.Y. lab for discussion and Claire Delahunty for critically perusing the manuscript. We thank Saman Eskandarzadeh, Michael Kubitz, and Erik Georgeson in W.R.S. lab for assistance in protein production. This work was supported by the grants P41GM103533 (NIH) and UM1AI100663, UM1AI144462, R01/R56AI113867 (NIH/NIAID). We thank Bill and Melinda Gates Foundation (BMGF) Collaboration for AIDS Vaccine Discovery (CAVD) funding to IAVI Neutralizing Antibody Center (NAC) and Ragon Institute post-doctoral fellowship to T.S.

## Author Contributions

S.B. conceived the method. S.B., J.K.D. and X.W. designed the experiments. S.B. processed the samples and analyzed data. J.K.D. performed LC-MS/MS. S.M.B. created the GlycoMSQuant tool. T.S. and B.G. expressed and purified HIV *Env* trimers. W.R.S., J.C.P. and J.R.Y. supervised the project. S.B. wrote the paper with contribution from all authors.

## Data Availability

Mass spectrometry data has been deposited in MassIVE-KB repository and is also accessible through ProteomeXchange Consortium with identifiers MSV000087414 and PXD025990, respectively.

## Code Availability

GlycoMSQuant source code is freely available at https://github.com/proteomicsyates/GlycoMSQuant under a permissive Apache License 2.0.

## References

1 Rudd, P. M. & Dwek, R. A. Glycosylation: heterogeneity and the 3D structure of proteins. Crit Rev Biochem Mol Biol 32, 1–100, doi:10.3109/10409239709085144 (1997).

2 Burton, D. R. & Hangartner, L. Broadly Neutralizing Antibodies to HIV and Their Role in Vaccine Design. Annu Rev Immunol 34, 635–659, doi:10.1146/annurev-immunol-041015-055515 (2016).

3 Cao, L. et al. Global site-specific N-glycosylation analysis of HIV envelope glycoprotein. Nat Commun 8, 14954, doi:10.1038/ncomms14954 (2017).

4 MacCoss, M. J. et al. Shotgun identification of protein modifications from protein complexes and lens tissue. Proc Natl Acad Sci U S A 99, 7900–7905, doi:10.1073/pnas.122231399 (2002).

5 Zhang, Y., Fonslow, B. R., Shan, B., Baek, M. C. & Yates, J. R., 3rd. Protein analysis by shotgun/bottom-up proteomics. Chem Rev 113, 2343–2394, doi:10.1021/cr3003533 (2013).

6 Tsiatsiani, L. & Heck, A. J. Proteomics beyond trypsin. Febs j 282, 2612–2626, doi:10.1111/febs.13287 (2015).

7 Behrens, A. J. et al. Molecular Architecture of the Cleavage-Dependent Mannose Patch on a Soluble HIV-1 Envelope Glycoprotein Trimer. J Virol 91, doi:10.1128/jvi.01894-16 (2017).

8 Go, E. P. et al. Glycosylation Benchmark Profile for HIV-1 Envelope Glycoprotein Production Based on Eleven Env Trimers. J Virol 91, doi:10.1128/jvi.02428-16 (2017).

9 Wu, C. C., MacCoss, M. J., Howell, K. E. & Yates, J. R., 3rd. A method for the comprehensive proteomic analysis of membrane proteins. Nat Biotechnol 21, 532–538, doi:10.1038/nbt819 (2003).

10 Steichen, J. M. et al. HIV Vaccine Design to Target Germline Precursors of Glycan-Dependent Broadly Neutralizing Antibodies. Immunity 45, 483–496, doi:10.1016/j.immuni.2016.08.016 (2016).

11 Steichen, J. M. et al. A generalized HIV vaccine design strategy for priming of broadly neutralizing antibody responses. Science 366, doi:10.1126/science.aax4380 (2019).

12 Sanders, R. W. et al. A next-generation cleaved, soluble HIV-1 Env trimer, BG505 SOSIP.664 gp140, expresses multiple epitopes for broadly neutralizing but not non-neutralizing antibodies. PLoS Pathog 9, e1003618, doi:10.1371/journal.ppat.1003618 (2013).

13 Meier, F. et al. Online Parallel Accumulation-Serial Fragmentation (PASEF) with a Novel Trapped Ion Mobility Mass Spectrometer. Mol Cell Proteomics 17, 2534–2545, doi:10.1074/mcp.TIR118.000900 (2018).

14 Seabright, G. E., Doores, K. J., Burton, D. R. & Crispin, M. Protein and Glycan Mimicry in HIV Vaccine Design. J Mol Biol 431, 2223–2247, doi:10.1016/j.jmb.2019.04.016 (2019).

15 Zhou, H. X., Rivas, G. & Minton, A. P. Macromolecular crowding and confinement: biochemical, biophysical, and potential physiological consequences. Annu Rev Biophys 37, 375–397, doi:10.1146/annurev.biophys.37.032807.125817 (2008).

16 Bar-On, Y. M., Flamholz, A., Phillips, R. & Milo, R. SARS-CoV-2 (COVID-19) by the numbers. Elife 9, doi:10.7554/eLife.57309 (2020).

17 Scanlan, C. N. et al. Inhibition of mammalian glycan biosynthesis produces non-self antigens for a broadly neutralising, HIV-1 specific antibody. J Mol Biol 372, 16–22, doi:10.1016/j.jmb.2007.06.027 (2007).

18 Fan, J. Q. & Lee, Y. C. Detailed studies on substrate structure requirements of glycoamidases A and F. J Biol Chem 272, 27058–27064, doi:10.1074/jbc.272.43.27058 (1997).

19 He, L., Diedrich, J., Chu, Y. Y. & Yates, J. R., 3rd. Extracting Accurate Precursor Information for Tandem Mass Spectra by RawConverter. Anal Chem 87, 11361–11367, doi:10.1021/acs.analchem.5b02721 (2015).

20 Xu, T. et al. ProLuCID: An improved SEQUEST-like algorithm with enhanced sensitivity and specificity. J Proteomics 129, 16–24, doi:10.1016/j.jprot.2015.07.001 (2015).

21 Tabb, D. L., McDonald, W. H. & Yates, J. R., 3rd. DTASelect and Contrast: tools for assembling and comparing protein identifications from shotgun proteomics. J Proteome Res 1, 21–26, doi:10.1021/pr015504q (2002).

22 Peng, J., Elias, J. E., Thoreen, C. C., Licklider, L. J. & Gygi, S. P. Evaluation of multidimensional chromatography coupled with tandem mass spectrometry (LC/LC-MS/MS) for large-scale protein analysis: the yeast proteome. J Proteome Res 2, 43–50, doi:10.1021/pr025556v (2003).

23 Park, S. K., Venable, J. D., Xu, T. & Yates, J. R., 3rd. A quantitative analysis software tool for mass spectrometry-based proteomics. Nat Methods 5, 319–322, doi:10.1038/nmeth.1195 (2008).

24 Zhang, M. et al. Tracking global patterns of N-linked glycosylation site variation in highly variable viral glycoproteins: HIV, SIV, and HCV envelopes and influenza hemagglutinin. Glycobiology 14, 1229–1246, doi:10.1093/glycob/cwh106 (2004).

25 Mann, H. B. & Whitney, D. R. On a Test of Whether one of Two Random Variables is Stochastically Larger than the Other. The Annals of Mathematical Statistics 18, 50–60 (1947).

26 Benjamini, Y. & Hochberg, Y. Controlling the False Discovery Rate: A Practical and Powerful Approach to Multiple Testing. Journal of the Royal Statistical Society. Series B (Methodological) 57, 289–300 (1995).

