## Supplementary Figures for "DeGlyPHER: an ultrasensitive method for analysis of viral spike N-glycoforms"

**Department of Molecular Medicine, The Scripps Research Institute, La Jolla, CA, USA**

Sabyasachi Baboo, Jolene K Diedrich, Salvador Martínez-Bartolomé, Xiaoning Wang, James C Paulson and John R Yates III

**Department of Immunology and Microbiology, The Scripps Research Institute, La Jolla, CA, USA**

Torben Schiffner, Bettina Groschel, William R Schief, James C Paulson

**IAVI Neutralizing Antibody Center, The Scripps Research Institute, La Jolla, CA, USA**

Torben Schiffner, Bettina Groschel, William R Schief

**The Ragon Institute of Massachusetts General Hospital, Massachusetts Institute of Technology and Harvard, Cambridge, MA, USA**

Torben Schiffner, William R Schief

**Corresponding authors:**

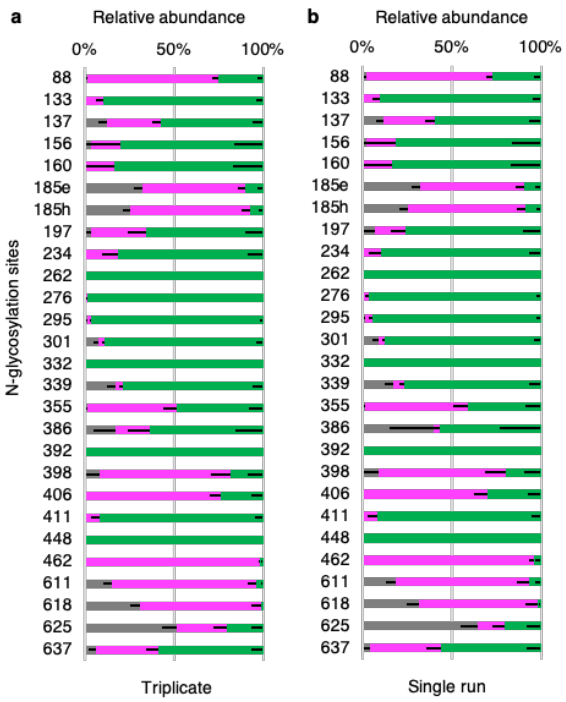


**Supplementary Figure 1 l N-glycan microheterogeneity in BG505 SOSIP.664 MD39 trimer using QE-HFX/nLC.** (**a**) Pattern observed using 4 h triplicates of 0.5 $\mu$g each from 5 $\mu$g starting material (**b**) A single run from the triplicate represents the same pattern (**c**) Trends at individual NGS remain mostly similar when triplicates of starting material ranging from 1 $\mu$g to 5 ng are processed. (**d**) Peptides identified per NGS across the dilution series. A change in N-glycan microheterogeneity at any NGS is attributed to low peptide sampling at that NGS due to lower starting material. N-glycosylation states are color-coded. Error bars represent mean-SEM. Between (**a**) and (**b**), there was no significant difference (BH-corrected $p$-value < 0.5) found in the proportion of any N-glycosylation state at any NGS.

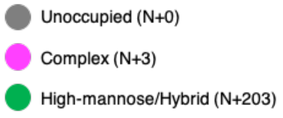


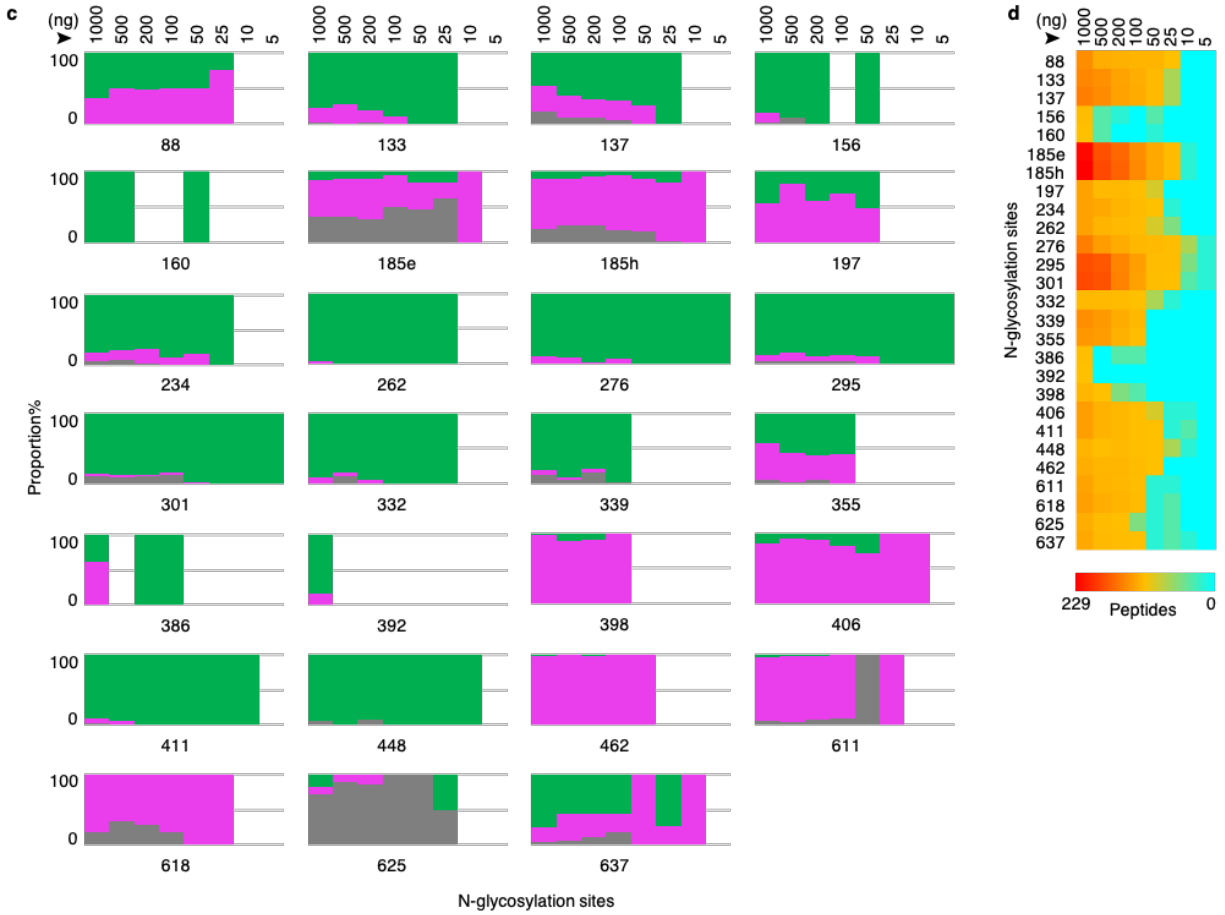

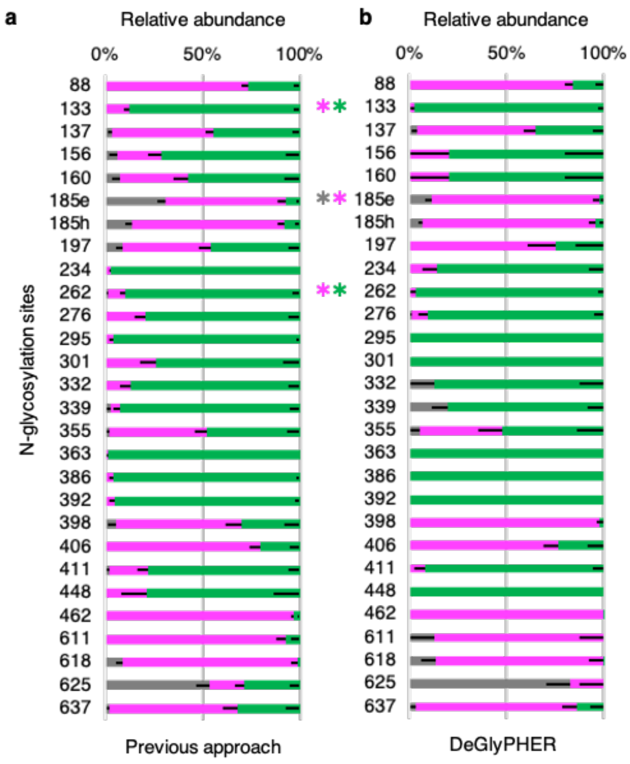


**Supplementary Figure 2 | Comparing methods using BG505 SOSIP.664 trimer.** The N-glycan microheterogeneity pattern is similar when comparing **(a)** our previous approach that used 18-fold more starting material and 9-fold more instrument time, when compared to **(b)** DeGlyPHER, which can reproduce the results with a single LC-MS/MS run, using a significantly shorter and simpler workflow. For **(b)**, data is from single 4 h runs of 0.5 $\mu$g from 5 $\mu$g starting material, using QE-HFX/nLC. N-glycosylation states are color-coded. Error bars represent mean-SEM. Between **(a)** and **(b)**, any significant difference (BH-corrected *p*-value <0.05) in proportion of a certain N-glycosylation state at any NGS is represented by color-coded ✱, and can be attributed to up to 3-fold higher peptides in (**a**) than (**b**), thus even small differences are found to be significant.

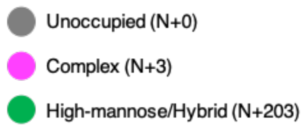

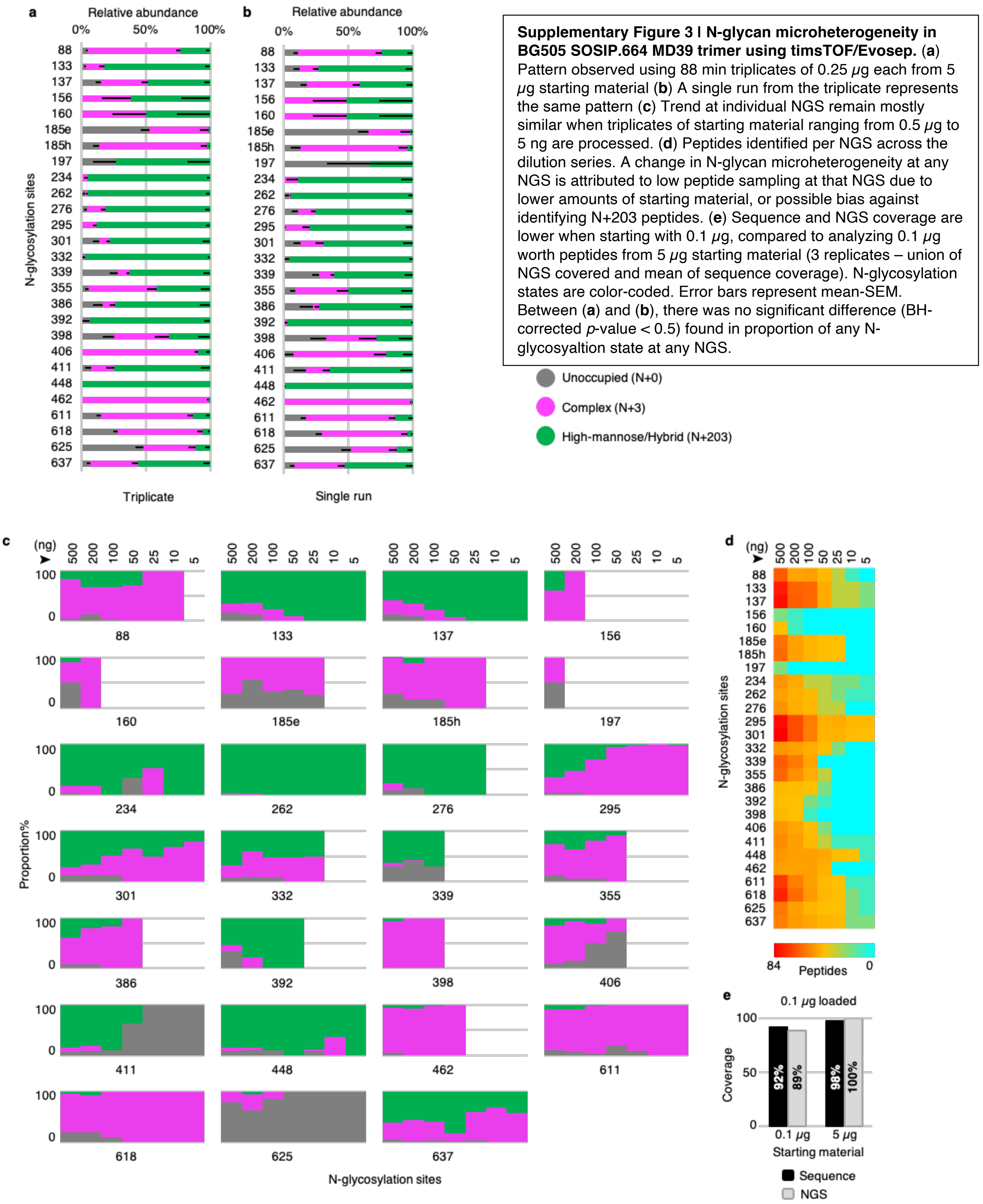


**Supplementary Figure 3 I N-glycan microheterogeneity in BG505 SOSIP.664 MD39 trimer using timsTOF/Evosep.** (**a**) Pattern observed using 88 min triplicates of 0.25 $\mu$g each from 5 $\mu$g starting material (**b**) A single run from the triplicate represents the same pattern (**c**) Trend at individual NGS remain mostly similar when triplicates of starting material ranging from 0.5 $\mu$g to 5 ng are processed. (**d**) Peptides identified per NGS across the dilution series. A change in N-glycan microheterogeneity at any NGS is attributed to low peptide sampling at that NGS due to lower amounts of starting material, or possible bias against identifying N+203 peptides. (**e**) Sequence and NGS coverage are lower when starting with 0.1 $\mu$g, compared to analyzing 0.1 $\mu$g worth peptides from 5 $\mu$g starting material (3 replicates – union of NGS covered and mean of sequence coverage). N-glycosylation states are color-coded. Error bars represent mean-SEM. Between (**a**) and (**b**), there was no significant difference (BH-corrected $p$-value < 0.5) found in proportion of any N-glycosyaltion state at any NGS.

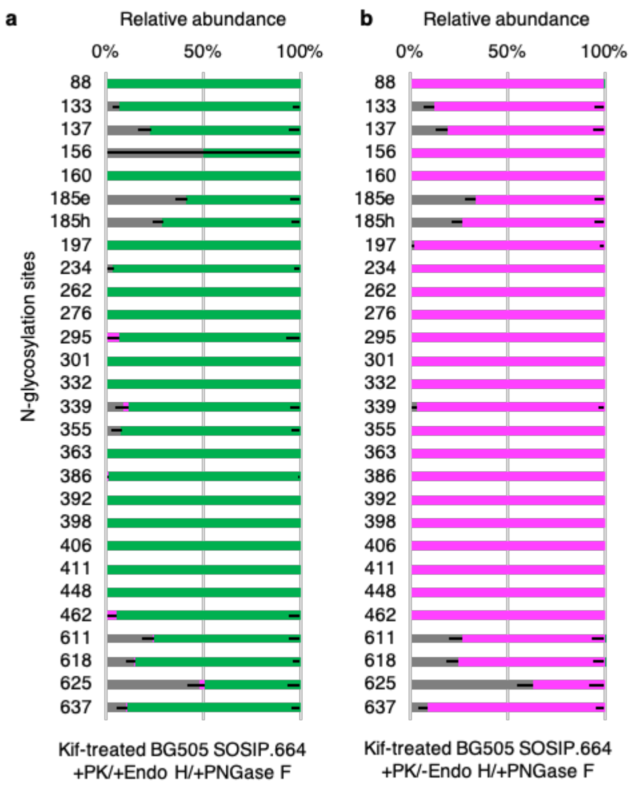


**Supplementary Figure 4 I Evaluating efficiency of glycosidases using Kifunensine-treated BG505 SOSIP.664 trimer.** (**a**) As expected, when treated with Endo H, followed by PNGase F, all NGS show predominantly +203 mass modification. (**b**) In the absence of Endo H treatment, and when treated with PNGase F, all NGS show predominantly +3 mass modification. Data is from single 4 h runs of 0.5 $\mu$g from 5 $\mu$g starting material, using QE-HFX/nLC. N-glycosylation states are color-coded. Error bars represent mean-SEM.

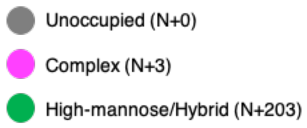

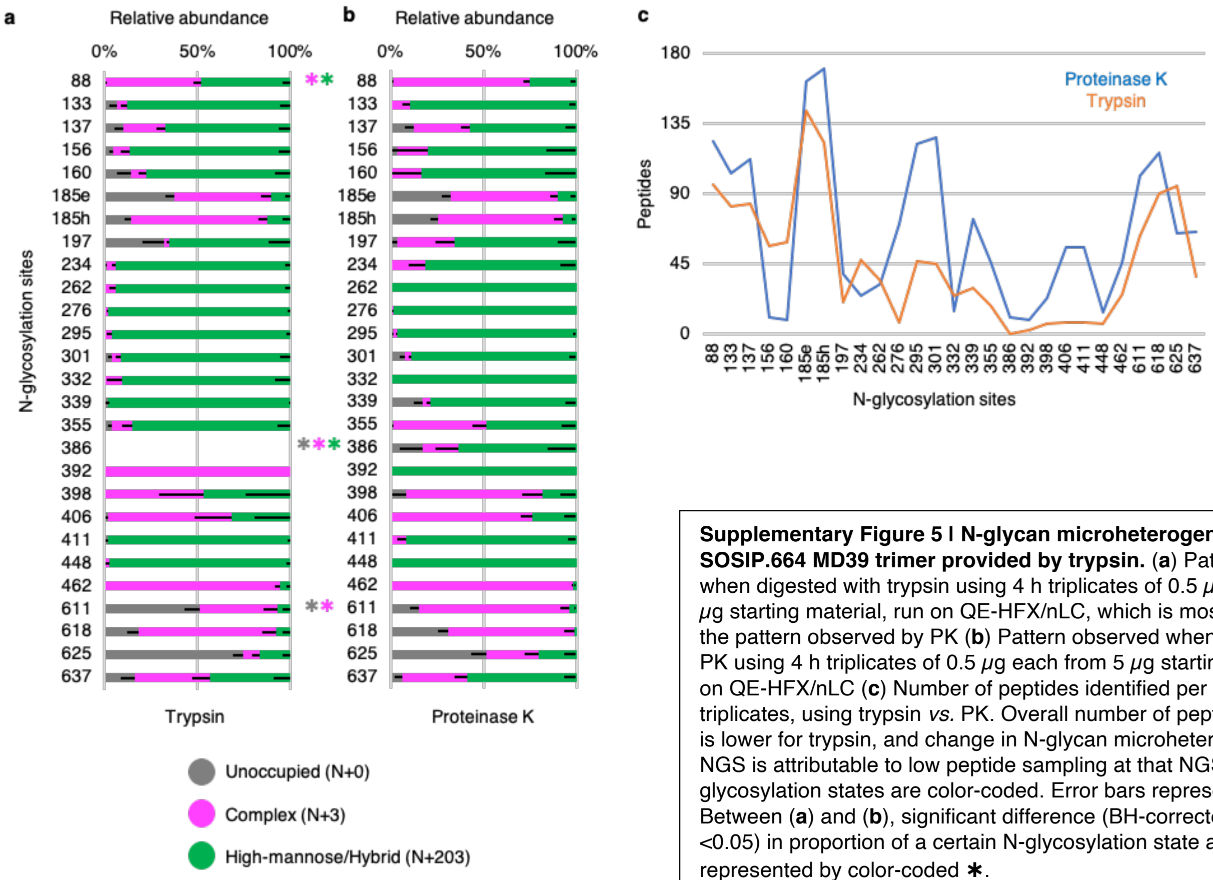


**Supplementary Figure 5 I N-glycan microheterogeneity in BG505 SOSIP.664 MD39 trimer provided by trypsin.** (**a**) Pattern observed when digested with trypsin using 4 h triplicates of 0.5 $\mu$g each from 5 $\mu$g starting material, run on QE-HFX/nLC, which is mostly similar to the pattern observed by PK (**b**) Pattern observed when digested with PK using 4 h triplicates of 0.5 $\mu$g each from 5 $\mu$g starting material, run on QE-HFX/nLC (**c**) Number of peptides identified per NGS across triplicates, using trypsin *vs.* PK. Overall number of peptides identified is lower for trypsin, and change in N-glycan microheterogeneity at any NGS is attributable to low peptide sampling at that NGS. N-glycosylation states are color-coded. Error bars represent mean-SEM. Between (**a**) and (**b**), significant difference (BH-corrected *p*-value <0.05) in proportion of a certain N-glycosylation state at any NGS is represented by color-coded ✱.
